# Gene Expression Imputation with Generative Adversarial Imputation Nets

**DOI:** 10.1101/2020.06.09.141689

**Authors:** Ramon Viñas, Tiago Azevedo, Eric R. Gamazon, Pietro Liò

## Abstract

A question of fundamental biological significance is to what extent the expression of a subset of genes can be used to recover the full transcriptome, with important implications for biological discovery and clinical application. To address this challenge, we present GAIN-GTEx, a method for gene expression imputation based on Generative Adversarial Imputation Networks. In order to increase the applicability of our approach, we leverage data from GTEx v8, a reference resource that has generated a comprehensive collection of transcriptomes from a diverse set of human tissues. We compare our model to several standard and state-of-the-art imputation methods and show that GAIN-GTEx is significantly superior in terms of predictive performance and runtime. Furthermore, our results indicate strong generalisation on RNA-Seq data from 3 cancer types across varying levels of missingness. Our work can facilitate a cost-effective integration of large-scale RNA biorepositories into genomic studies of disease, with high applicability across diverse tissue types.

## 1 Introduction

High-throughput profiling of the transcriptome has revolutionised discovery methods in the biological sciences. The resulting gene expression measurements can be used to uncover disease mechanisms (Cookson et al., 2009; Emilsson et al., 2008; Gamazon et al., 2018), propose novel drug targets (Evans and Relling, 2004; Sirota et al., 2011), provide a basis for comparative genomics (Colbran et al., 2019; King and Wilson, 1975), and motivate a wide range of fundamental biological problems.

One question of fundamental biological significance is to what extent the expression of a subset of genes can be used to recover the full transcriptome with minimal reconstruction error. Genes that participate in similar biological processes or that have shared molecular function are likely to have similar expression profiles (Zhang and Horvath, 2005), raising the possibility of gene expression prediction from a minimal subset of genes. Moreover, gene expression measurements may suffer from unreliable values because some regions of the genome are extremely challenging to interrogate due to high genomic complexity or sequence homology (Conesa et al., 2016), highlighting the need for accurate imputation. Most gene expression studies continue to be performed with specimens derived from peripheral blood or a surrogate (e.g. lymphoblastoid cell lines; LCLs) due to the difficulty of collecting some tissues. However, gene expression may be highly tissue or cell-type specific, potentially limiting the utility of a proxy tissue.

The missing data problem can adversely affect downstream gene expression analysis. The simple approach of excluding samples with missing data from the analysis can lead to a substantial loss in statistical power. Dimensionality reduction approaches such as principal component analysis (PCA) and singular value decomposition (SVD) (Wall et al., 2003) cannot be applied to gene expression data with missing values. Clustering methods, a mainstay of genomics, such as k-means and hierarchical clustering may become unstable even with a few missing values (Troyanskaya et al., 2001).

To address these challenges, we develop an approach to gene expression imputation based on Generative Adversarial Imputation Nets (GAIN; Yoon et al., 2018), which we name GAIN-GTEx. We present an architecture that recovers missing expression data for multiple tissue types under the *missing completely at random* assumption (MCAR; Little and Rubin, 2019), that is, the missingness of the data is independent of the observed and the missing variables. In contrast to existing linear methods for deconfounding gene expression (Øystein Sørensen et al., 2018), our model integrates covariates (global determinants of gene expression; Stegle et al., 2012) to account for their non-linear effects. In particular, a characteristic feature of our architecture is the use of word embeddings (Mikolov et al., 2013) to learn rich and distributed representations for the tissue types and other covariates. To enlarge the possibility and scale of a study’s expression data (e.g. by including samples from highly inaccessible tissues), we train our model on RNA-Seq data from the Genotype-Tissue Expression (GTEx) project (Consortium et al., 2015; Consortium et al., 2017), a reference resource (v8) that has generated a comprehensive collection of human transcriptome data in a diverse set of tissues.

We show that our method is superior to several standard and state-of-the-art imputation methods in terms of predictive performance and runtime. Furthermore, we demonstrate that our model is highly applicable across many different tissues and varying levels of missingness. Finally, to analyse the cross-study relevance of our method, we evaluate GAIN-GTEx on gene expression data from The Cancer Genome Atlas (TCGA; Weinstein et al., 2013a) and show that our approach is robust when applied to RNA-Seq data from 3 different cancers. Our code can be found at: https://github.com/rvinas/GAIN-GTEx.

## 2 Methods

In this section, we introduce our approach to imputing gene expression. Throughout the remainder of the paper, we use script letters to denote sets (e.g. 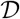), upper-case bold symbols to denote matrices or random variables (e.g. **X**), and lower-case bold symbols to denote column vectors (e.g. **x** or 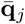). The rest of the symbols (e.g. 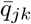, *G*, or *f*) denote scalar values or functions.

### 2.1 Problem formulation

Consider a dataset 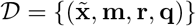, where 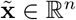 represents a vector of gene expression values with missing components; **m** ∈ {0,1}^n^ is a mask indicating which components of the original vector of expression values **x** are missing or observed; *n* is the number of genes; and 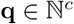 and 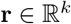 are vectors of c categorical (e.g. tissue type or sex) and *k* quantitative covariates (e.g. age), respectively. Our goal is to recover the original gene expression vector 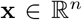 by modelling the conditional probability distribution 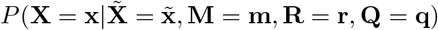. By modelling the distribution, we can also quantify the uncertainty of the imputed expression values.

### 2.2 GAIN for gene expression imputation

Our method builds on Generative Adversarial Imputation Nets (GAIN; Yoon et al., 2018). Similar to generative adversarial networks (GANs; Goodfellow et al., 2014), GAIN estimates a generative model via an adversarial process driven by the competition between two players, the *generator* and the *discriminator*.

#### Generator

The generator aims at recovering missing data from partial gene expression observations, producing samples from the conditional 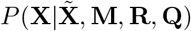. Formally, we define the generator as a function 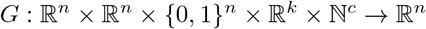 that imputes expression values as follows:

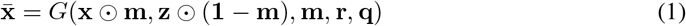

where 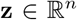 is a vector sampled from a fixed noise distribution. Here ʘ denotes element-wise multiplication. Similar to GAIN, we mask the n-dimensional noise vector as **z** ʘ (**1** − **m**), encouraging a bijective association between noise components and genes. Before passing the output 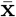 to the discriminator, we replace the prediction for the non-missing components by the original, observed expression values:

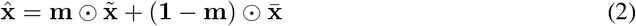

#### Discriminator

The discriminator takes the imputed samples 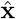 and attempts to distinguish whether the expression value of each gene has been observed or produced by the generator. This is in contrast to the original GAN discriminator, which receives information from two input streams (generator and data distribution) and attempts to distinguish the true input source.

Formally, the discriminator is a function 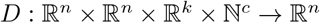 that outputs the probability 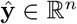 of each gene being observed as opposed to being imputed by the generator:

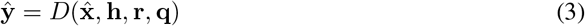

Here, the vector 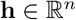 corresponds to the *hint* mechanism described in Yoon et al. (2018), which provides theoretical guarantees on the uniqueness of the global minimum for the estimation of 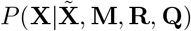. Concretely, the role of the hint vector **h** is to *leak* some information about the mask **m** to the discriminator. Similar to GAIN, we define the hint **h** as follows:

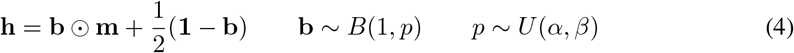

where **b** ∈ {0,1}^*n*^ is a binary vector that controls the amount of information from the mask **m** revealed to the discriminator. In contrast to GAIN, which discloses all but one components of the mask, we sample **b** from a Bernoulli distribution parametrised by a random probability *p ~ U*(*α, β*), where *α* ∈ [0,1] and *β* ∈ [α, 1] are hyperparameters. This accounts for a high number of genes n and allows to trade off the number of mask components that are revealed to the discriminator.

#### Optimisation

Similarly to GAN and GAIN, we optimise the generator and discriminator adversarially, interleaving gradient updates for the discriminator and generator.

The discriminator aims at determining whether genes have been observed or imputed based on the imputed vector 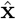, the covariates **q** and **r**, and the hint vector **h**. Since the hint vector **h** readily provides partial information about the ground truth **m** (Equation 4), we penalise D only for genes *i* ∈ {1,2,..., *n*} such that *h_i_* = 0.5, that is, genes whose corresponding mask value is unavailable to the discriminator. We achieve this via the following loss function 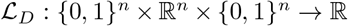:

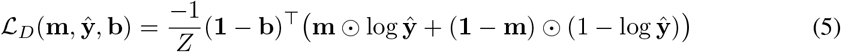

where *Z* = 1 + (**1** − **b**)^⊤^ (**1** − **b**) is a normalisation term. The only difference with respect to the binary cross entropy loss function is the dot product involving (**1** − **b**), which we employ to ignore genes whose mask has been *leaked* to the discriminator through **h**.

The generator aims at implicitly estimating 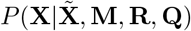. Therefore, its role is not only to impute the expression corresponding to missing genes, but also to reconstruct the expression of the observed inputs. Similar to GAIN, in order to account for this and encourage a realistic imputation function, we use the following loss function 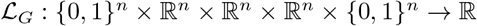 for the generator:

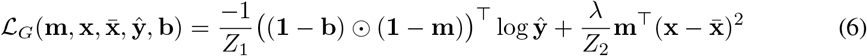

where *Z*_1_ = 1 + (**1** − **b**)^⊤^(**1** − **b**) and *Z*_2_ = **m**^⊤^**m** are normalisation terms, and λ > 0 is a hyperparameter. Intuitively, the first term in Equation 6 corresponds to the adversarial loss, whereas the second term accounts for the loss that the generator incurs in the reconstruction of the observed gene expression values.

#### Architecture

We model the discriminator D and the generator G using neural networks. We first describe how we use word embeddings, a distinctive feature of both models that allow to learn rich, dense representations for the different tissue types and, more generally, for all the covariates 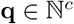.

Formally, let *q_j_* be a categorical covariate (e.g. tissue type) with vocabulary size *v_j_*, that is, *q_j_* ∈ {1,2,..., v_j_}, where each value in the vocabulary {1,2,..., v_j_} represents a different category (e.g. whole blood or kidney). Let 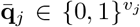 be a one-hot vector such that 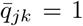 if *q_j_* = *k* and 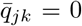 otherwise. Let *d_j_* be the dimensionality of the embeddings for covariate *j*. We obtain a vector of embeddings 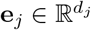 as follows:

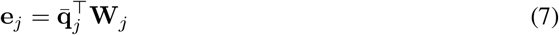

where each 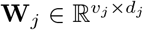 is a matrix of learnable weights. Essentially, this operation describes a lookup search in a dictionary with *v_j_* entries, where each entry contains a learnable *d_j_*-dimensional vector of embeddings that characterise each of the possible values that *q_j_* can take. To obtain a global collection of embeddings **e**, we concatenate all the vectors e_*j*_ for each categorical covariate *j*:

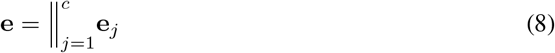

where *c* is the number of categorical covariates and ≑ represents the concatenation operator. We then use the learnable embeddings e in downstream tasks.

In terms of the architecture, we model both the generator *G* and discriminator *D* as neural networks that leverage independent instances **e**^*G*^ and **e**^*D*^ of the categorical embeddings for their corresponding downstream tasks. Specifically, we model the two players as follows:

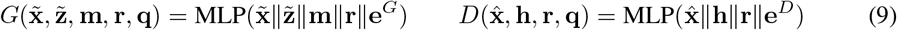

where MLP denotes a multilayer perceptron 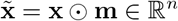 and 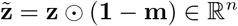 are the masked gene expression and noise input vectors, respectively. Figure 1 shows the architecture of both players.

**Figure 1:**
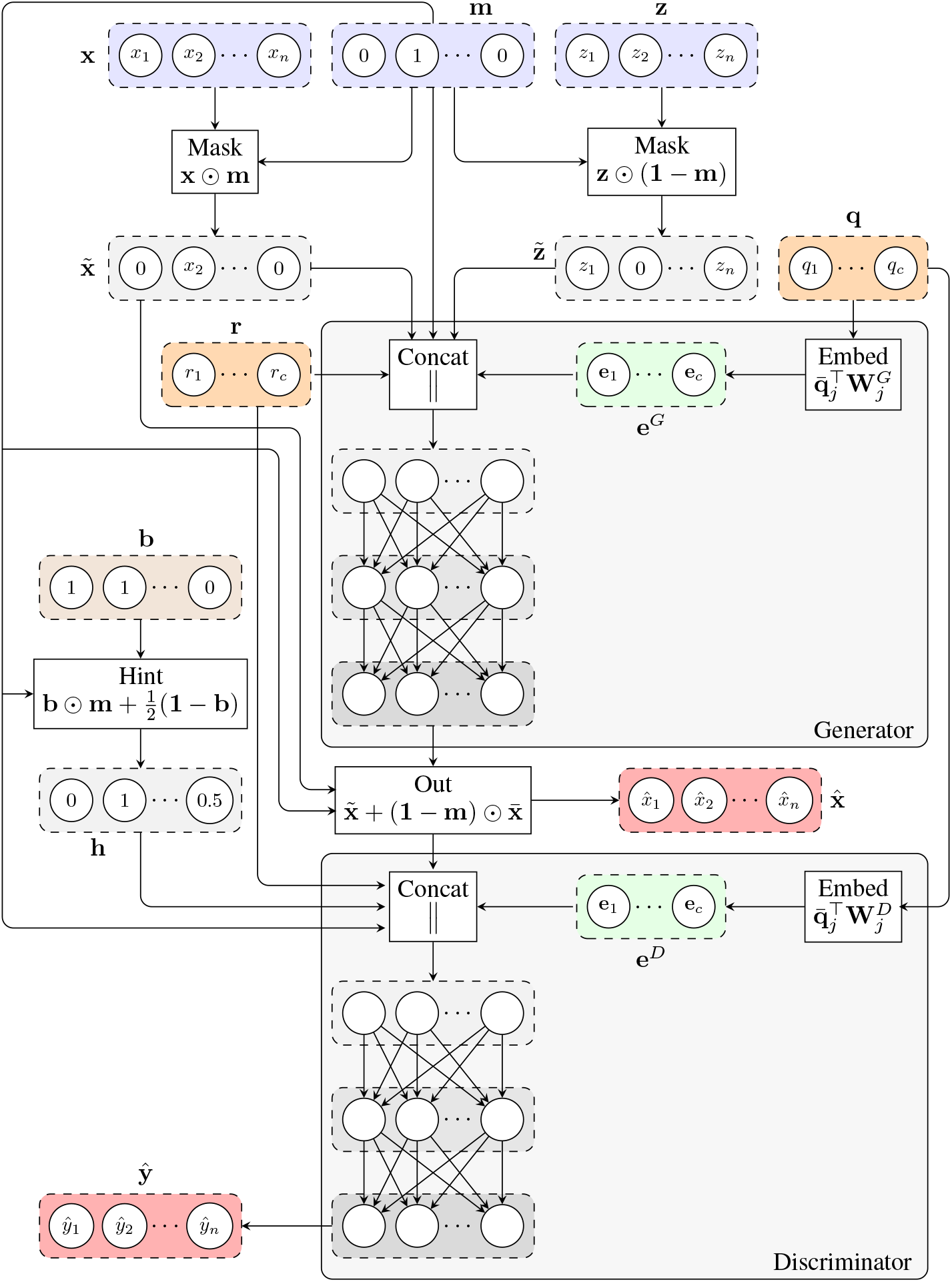
Architecture of GAIN-GTEx. The generator takes gene expression values 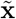 with missing components according to a mask **m**, and categorical (e.g. tissue type; **q**) and numerical (e.g. age; **r**) covariates, and outputs the imputed values 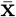. The observed components of the generator’s output are then replaced by the actual observed expression values 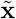, yielding the imputed sample 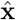. The discriminator receives 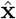, the sample covariates, and the hint vector **h**, and produces the probabilities 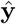 of each gene being observed as opposed to being imputed by the generator.

## 3 Results

Here, we evaluate the performance of the proposed model on the GTEx dataset (Aguet et al., 2019). In the first section, we provide an overview of the dataset and describe the experimental details, including the hyperparameters of our model and how we generate the masks for training and evaluation. In the second section, we present the imputation results and compare our method to standard and state-of-the-art imputation algorithms for gene expression data.

### 3.1 Experimental details

#### Dataset

The GTEx dataset is a public genomic resource of genetic effects on the transcriptome across a broad collection of human tissues, enabling linking of these regulatory mechanisms to trait and disease associations (Aguet et al., 2019). Our dataset contains 15,201 RNA-Seq samples collected from 49 tissues of 838 unique donors. We also select the intersection of all the protein-coding genes among these tissues, yielding 12,557 unique human genes. In addition to the expression data, we leverage metadata about the sample donors^1^, including sex, age, and cohort (post-mortem, surgical or organ donor). We normalise the expression data via the standard score, so that the standardised expression values have mean 0 and standard deviation 1 for each gene across all samples.

To prevent any leakage of information between the train and test sets, we enforce all samples from the same donor to be within the same set. Concretely, we first flip the GTEx donor identifiers (e.g. 111CU-1826 is flipped to 6281-UC111), we then sort the reversed identifiers in alphabetical order, and we finally select a suitable split point, forcing the two sets to be disjoint. After splitting the data, the train set, which we use to train and tune the model, consists of ~ 75% of the total samples, while the test set, on which we evaluate the final performance, contains the remaining data.

#### GAIN implementation

For each sample, we include the donor’s age as numerical covariate in **r** and the tissue type, sex and cohort as categorical covariates in **q**. We normalise the numerical variables via the standard score. For each categorical variable *q_j_* ∈ {1,2,..., *v_j_*}, we use the rule of thumb 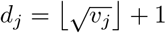 to set all the dimensions of the categorical embeddings for both players.

In terms of the MLP architectures, we use two hidden layers for both players, with 256 units and ReLU activation functions each. Regarding the output activation, we leverage a linear and a sigmoid activation functions for the generator and discriminator, respectively. The linear activation ensures that the range of the output expression is unrestricted. We find that adding more hidden layers in the generator or discriminator networks does not yield significant improvements in our validation scores.

We train our model using RMSProp (Tieleman and Hinton, 2012) with a learning rate of 0.001. We use batch normalisation (Ioffe and Szegedy, 2015) in the hidden layers of both the generator and discriminator, which yields a significant speed-up to the training convergence according to our experiments. Regarding the hyperparameter λ to trade off the adversarial and reconstruction losses of the generator (see Equation 6), we find that setting λ = 1 yields good results. We use early stopping to train the model, stopping when the validation loss has not improved in the last 10 epochs. We train the model for ~ 9 hours on a NVIDIA TITAN XP GPU with 12 GB of RAM.

#### Mask and hint generation

We assume that the data is *missing completely at random* (MCAR), that is, the missingness does not depend on any of the observed nor unobserved variables and the conditional distribution *P*(**M**|**X**, **R**, **Q**) can be expressed as *P*(**M**). At training time, for each training example, we sample the mask vector **m** from a Bernoulli distribution *B*(1,*p*) parametrised by a random probability *p* ~ *U*(0.5,0.95). To generate the hint vector **h** (see Equation 4), we sample **b** from *B*(1,*p*), where *p* ~ *U*(0.8,0.95). At prediction time, the mask **m** ∈ {0,1}^n^ can be fixed to any desired value - for evaluation purposes we sample it from *B*(1,0.5) unless otherwise stated.

### 3.2 Imputation results

Here we provide an overview of the imputation results, including a comparison with other imputation methods, an evaluation of the tissue-specific results, and an analysis of the cross-study relevance of our method across different levels of missingness.

#### Baseline methods

First, we leverage two standard gene expression approaches as baselines: blood surrogate and median imputation. For blood surrogate imputation, we impute missing gene expression values in any given tissue with the corresponding expression values measured in whole blood for the same donor, as widely practised in biomedical research due to greater accessibility of blood-derived RNA sources. For median imputation, we impute missing values with the median of the observed tissue-specific gene expression computed across donors.

Second, we compare GAIN-GTEx to two state-of-the-art methods. The first, Multivariate Imputation by Chained Equations (MICE; Buuren and Groothuis-Oudshoorn, 2010), leverages chained equations to create multiple imputations of missing data. The second, MissForest (Stekhoven and Bühlmann, 2012), is a non-parametric imputation method based on random forests trained on observed values to impute the missing values.

Finally, we compare GAIN-GTEx to a simplified version which we call GAIN-MSE-GTEx. In contrast to the proposed method, GAIN-MSE-GTEx is trained exclusively on the mean squared error corresponding to the reconstruction loss of Equation 6, excluding the discriminator network. To train this model, we use the same generator architecture and hyperparameters as described in Section 3.1.

#### Comparison

Table 1 shows a quantitative summary of the imputation performances. In addition to the imputation scores, we include a *feasible* column that shows whether the gene expression imputation is computationally *feasible* under our experimental design. We label methods as computationally *unfeasible* when they take longer than 7 days to run on our server^2^, after which we halt the execution.

**Table 1:** Gene expression imputation performance with a missing rate of 50% across 10 runs. We do not report the *R*^2^ scores for MICE and MissForest, because the runtime is longer than 7 days, even for the original test set without oversampling.

| Method | $R^2$ | Feasible? |
| --- | --- | --- |
| MICE | — | × |
| MissForest | — | × |
| Blood surrogate | $-0.924 \pm 0.136$ | ✓ |
| Median imputation | $-0.024 \pm 0.012$ | ✓ |
| GAIN-MSE-GTEx | $0.649 \pm 0.021$ | ✓ |
| GAIN-GTEx | $0.659 \pm 0.022$ | ✓ |

In terms of the evaluation metrics, we report the coefficient of determination (*R*^2^). This metric ranges from −∞ to 1 and corresponds to the ratio of explained variance to the total variance. Notably, negative scores indicate that the model predictions are worse than those of a baseline model that predicts the mean of the data.

To evaluate the performance, we perform oversampling on the test set, obtaining an extended test set in which each tissue is equally represented. We then generate random masks with a missing rate of 50% and compute the imputation *R^2^* per gene. We repeat the last step 10 times and report the overall mean *R^2^* and the average per-gene standard deviation of the *R*^2^ scores, averaged across the 10 runs.

#### Tissue-specific results

Figure 2 shows the *R*^2^ scores achieved by GAIN-GTEx across 49 tissues. To obtain these results, we generate random masks with a missing rate of 50% for the extended test set, we perform imputation, and we plot the distribution of 12,557 gene *R*^2^ scores for each tissue.

**Figure 2:**
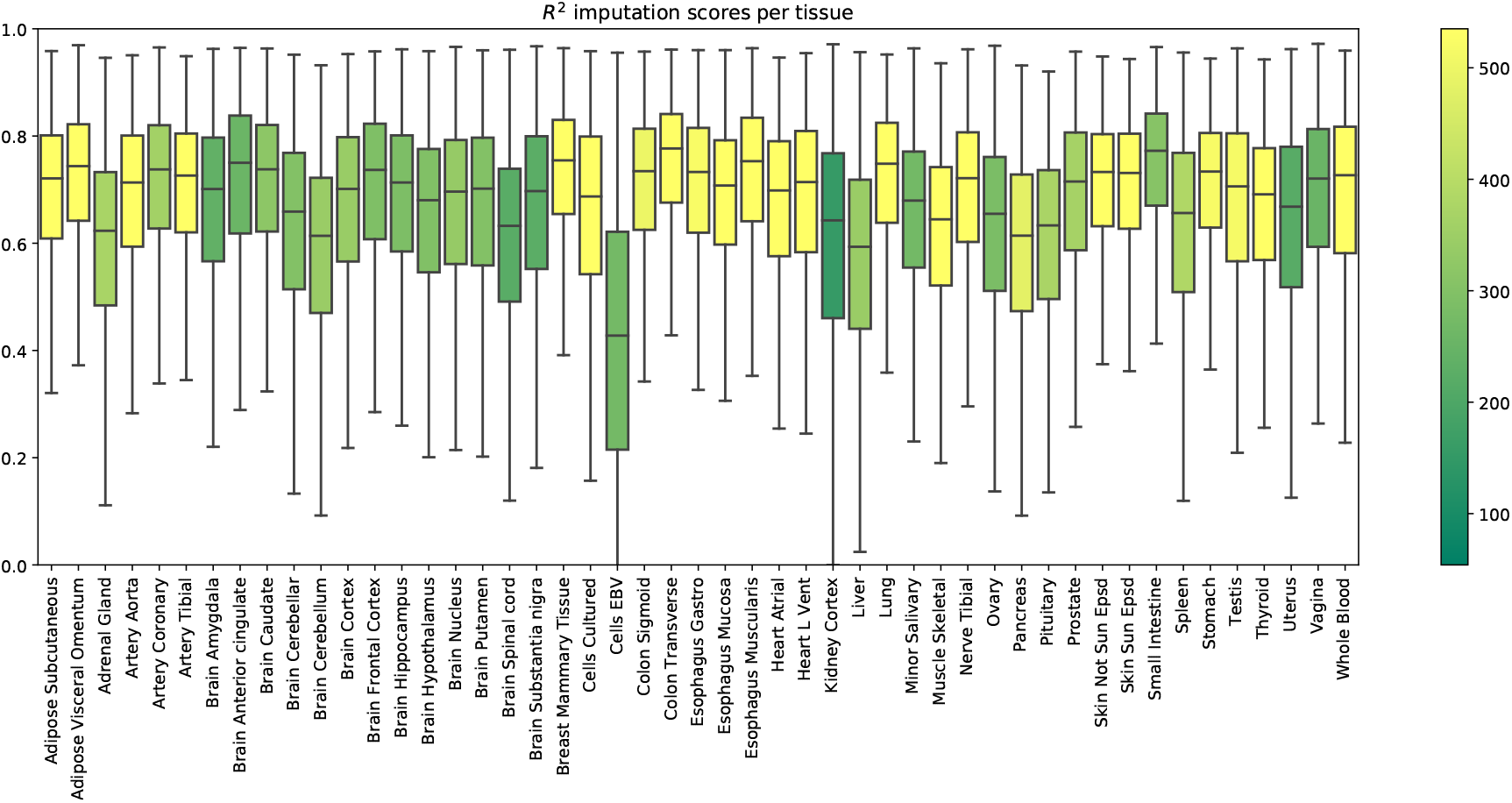
*R*^2^ imputation scores per GTEx tissue with a missing rate of 50%. Each box shows the distribution of the per-gene *R*^2^ scores in the extended test set. The colour of each box represents the number of training samples of the corresponding tissue.

In Figure 3, we illustrate the ability of GAIN-GTEx to learn rich tissue representations. Specifically, we plot a UMAP representation (McInnes et al., 2018) of the learnt tissue embeddings 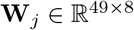 from the generator (see Equation 7), where *j* indexes the tissue dimension.

**Figure 3:**
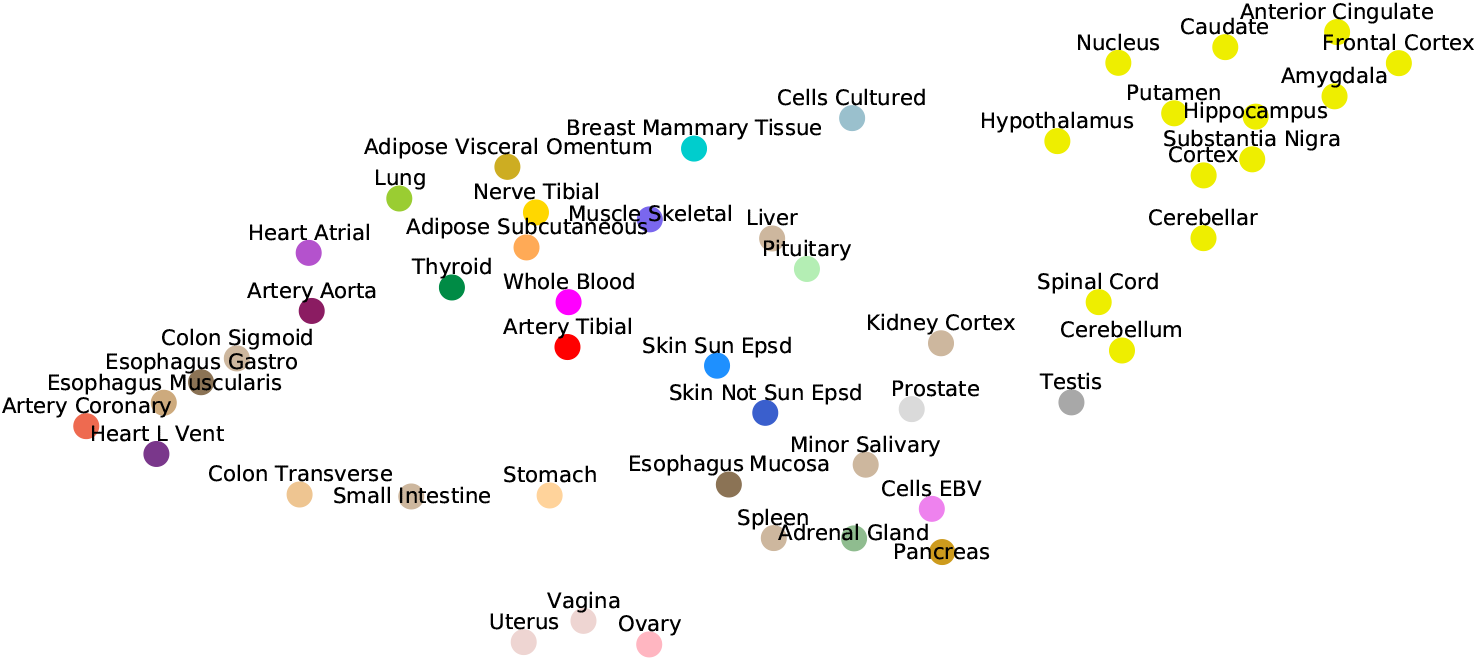
UMAP visualisation of the tissue embeddings from the generator. Colors are assigned to conform to the GTEx Consortium conventions. Note that the central nervous system, consisting of the 13 brain tissues, clusters together on the top right corner.

#### Cross-study results across missing rates

To evaluate the cross-study relevance of our method, we leverage the model trained on GTEx to perform imputation on The Cancer Genome Atlas (TCGA) gene expression data in acute myeloid leukemia (TCGA LAML; Network, 2013), breast cancer (TCGA BRCA; Network et al., 2012), and lung adenocarcinoma (TCGA LUAD; Network et al., 2014). For each TCGA tissue and its *non-diseased* test counterpart on GTEx, we show the imputation quality in Table 2 as well as the performance across varying missing rates in Figure 4.

**Table 2:** Cross-study results for our model trained on GTEx. For a missing rate of 50%, we report the *R*^2^ scores on data from 3 TCGA cancers and their *healthy* counterpart on GTEx (test set).

| Tissue | $R^2$ |
| --- | --- |
| TCGA LAML | $0.386 \pm 0.057$ |
| TCGA BRCA | $0.408 \pm 0.023$ |
| TCGA LUAD | $0.439 \pm 0.034$ |
| GTEx Whole blood | $0.678 \pm 0.031$ |
| GTEx Breast | $0.724 \pm 0.036$ |
| GTEx Lung | $0.713 \pm 0.033$ |

**Figure 4:**
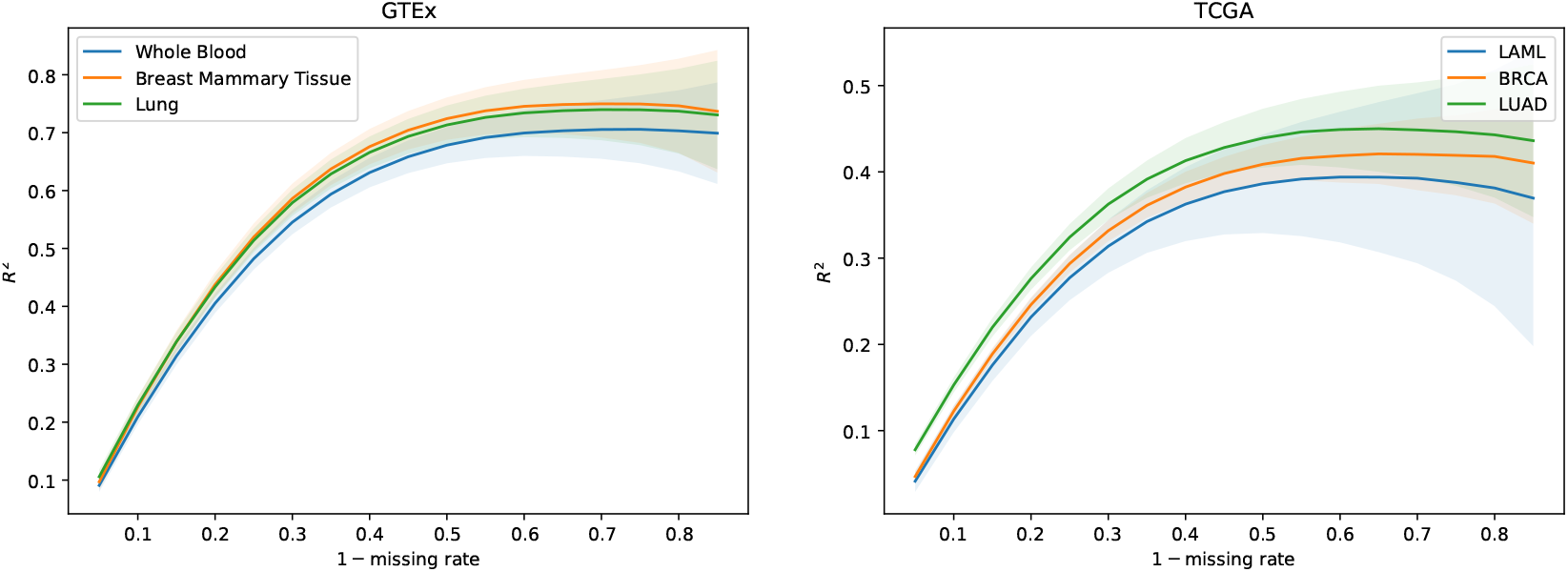
*R*^2^ imputation scores per tissue across missing rate for 3 TCGA cancers and their healthy counterpart in GTEx (test set). The shaded area represents one standard deviation of the per-gene *R*^2^ scores in the corresponding tissue. The greater the rate of missingness, the lower the performance.

## 4 Discussion

We develop an imputation approach to gene expression, facilitating the reconstruction of a high-dimensional molecular trait that is central to disease biology and drug target discovery. Our model builds on GAIN to learn complex probability distributions from incomplete gene expression data and relevant covariates (potential global determinants of expression). A characteristic feature of our architecture is the use of word embeddings in both players that enable them to learn distributed representations of the tissue types (see Figure 3).

To enlarge the possibility and scale of a study’s expression data, we leverage the most comprehensive human transcriptome resource available (GTEx), allowing us to test the performance of our method in a broad collection of tissues (see Figure 2). The biospecimen repository includes model systems such as whole blood and Epstein Barr virus (EBV) transformed lymphocytes; central nervous system tissues from 13 brain regions; and a wide diversity of other primary tissues from *non-diseased* individuals. In particular, we observe that EBV transformed lymphocytes, an accessible and renewable resource for functional genomics, are a notable outlier in imputation performance. This is perhaps not surprising, consistent with studies about the transcriptional effect of EBV infection on the suitability of the cell lines as a model system for primary tissues (Carter et al., 2002).

We compare our approach with several existing imputation methods and find that GAIN-GTEx significantly outperforms them in terms of imputation performance and runtime (see Table 1). We observe that standard approaches such as leveraging the expression of missing genes from a surrogate blood tissue yield negative *R*^2^ values and therefore do not perform well. Median imputation, although easy to implement, has a very limited predictive power. In terms of state-of-the-art-methods, we note that MICE and MissForest are computationally prohibitive given the high-dimensionality of the data and we halt the execution after running our experiments for 7 days. Finally, we observe that a simplification of our method, GAIN-MSE-GTEx, also performs well, suggesting that the mean squared error term of the generator’s loss function has a major role in the learning process.

To evaluate the cross-study relevance of our method, we apply the trained model derived from GTEx to perform imputation on The Cancer Genome Atlas gene expression data in acute myeloid leukemia, lung adenocarcinoma, and breast cancer. In addition to technical artifacts (e.g. batch effects), generalising to this data is highly challenging because the expression is largely driven by features of the disease such as chromosomal abnormalities, genomic instabilities, large-scale mutations, and epigenetic changes (Baylin and Jones, 2011; Weinstein et al., 2013b). Our results show that, despite these challenges, our method is robust to gene expression from multiple diseases in different tissues (see Table 2), lending itself to being used as a tool to extend independent transcriptomic studies. Finally, we evaluate the imputation performance of GAIN-GTEx for a range of values for the missing rate (see Figure 4). We note that the performance is stable and that the greater the proportion of missing values, the lower the prediction performance.

## 5 Conclusion

In this work, we develop a method for gene expression imputation based on generative adversarial imputation networks. To increase the applicability of our model, we train it on RNA-Seq data from the Genotype-Tissue Expression project, a reference resource that has generated a comprehensive collection of transcriptomes in a diverse set of tissues. A characteristic feature of the model is the use of word embeddings to learn distributed representations for the tissue types. Our approach achieves state-of-the-art performance in terms of both imputation quality and runtime, and generalises to transcriptomics data from 3 cancers of the The Cancer Genome Atlas. This work can facilitate the straightforward integration and cost-effective repurposing of large-scale RNA biorepositories into genomic studies of disease, with high applicability across diverse tissue types.

### Broader Impact

The study of the transcriptome is fundamental to our understanding of cellular and pathophysiological processes. High-dimensional gene expression data contain information relevant to a wide range of applications, including disease diagnosis (Huang et al., 2010), drug development (Sun et al., 2013), and evolutionary history (Colbran et al., 2019). Thus, accurate and robust methods for imputation of gene expression have the enormous potential to enhance our molecular understanding of complex diseases, inform the search for novel drugs, and provide key insights into evolutionary processes. Here, we develop a methodology that attains state-of-the art performance in terms of imputation quality and execution time. Our analysis shows that the use of blood as a surrogate for difficult-to-acquire tissues, as widely practised throughout biomedical research, has substantially degraded performance, with important implications for biomarker discovery and therapeutic development. Our method generalises to gene expression in a disease class which has shown considerable health outcome disparities across population groups in terms of morbidity and mortality. Future algorithmic developments therefore hold promise for more effective detection, diagnosis, and treatment (Hosny and Aerts, 2019) and for improved implementation in clinical medicine (Char et al., 2018). Increased availability of transcriptomes in diverse human populations to enlarge our training data (a well-known ethical challenge) should lead to further gains (i.e., decreased biases in results and reduced health disparities) (Wojcik et al., 2019). This work has the potential to catalyse research into the application of Generative Adversarial Networks (Goodfellow et al., 2014) for molecular reconstruction of cellular states and downstream gene mapping of complex traits (Cookson et al., 2009; Gamazon et al., 2015).

## Acknowledgments and Disclosure of Funding

We thank Nikola Simidjievski, Cătălina Cangea, Ben Day, Cristian Bodnar, and Arian Jamasb for the helpful comments and discussion. The project leading to these results has received funding from “la Caixa” Foundation (ID 100010434), under agreement LCF/BQ/EU19/11710059. This research is supported by the National Institutes of Health / NHGRIR35 HG010718 (ERG).

## Footnotes

1 The data and a detailed summary of the GTEx samples and donor information can be found at: https://gtexportal.org/home/tissueSummaryPage

2 CPU: Intel(R) Xeon(R) Processor E5-2630 v4. RAM: 125GB

## References

François Aguet, Alvaro N Barbeira, Rodrigo Bonazzola, Andrew Brown, Stephane E Castel, Brian Jo, Silva Kasela, Sarah Kim-Hellmuth, Yanyu Liang, Meritxell Oliva, Princy E Parsana, Elise Flynn, Laure Fresard, Eric R Gamazon, Andrew R Hamel, Yuan He, Farhad Hormozdiari, Pejman Mohammadi, Manuel Muñoz-Aguirre, YoSon Park, Ashis Saha, Ayellet V Segrć, Benjamin J Strober, Xiaoquan Wen, Valentin Wucher, Sayantan Das, Diego Garrido-Martín, Nicole R Gay, Robert E Handsaker, Paul J Hoffman, Seva Kashin, Alan Kwong, Xiao Li, Daniel MacArthur, John M Rouhana, Matthew Stephens, Ellen Todres, Ana Viñuela, Gao Wang, Yuxin Zou,, Christopher D Brown, Nancy Cox, Emmanouil Dermitzakis, Barbara E Engelhardt, Gad Getz, Roderic Guigo, Stephen B Montgomery, Barbara E Stranger, Hae Kyung Im, Alexis Battle, Kristin G Ardlie, and Tuuli Lappalainen. The gtex consortium atlas of genetic regulatory effects across human tissues. bioRxiv, 2019. doi: 10.1101/787903. URL https://www.biorxiv.org/content/early/2019/10/03/787903.

Stephen B Baylin and Peter A Jones. A decade of exploring the cancer epigenome—biological and translational implications. Nature Reviews Cancer, 11(10):726–734, 2011.

S van Buuren and Karin Groothuis-Oudshoorn. mice: Multivariate imputation by chained equations in r. Journal of statistical software, pages 1–68, 2010.

Kara L Carter, Ellen Cahir-McFarland, and Elliott Kieff. Epstein-barr virus-induced changes in b-lymphocyte gene expression. Journal of virology, 76(20):10427–10436, 2002.

Danton S Char, Nigam H Shah, and David Magnus. Implementing machine learning in health care—addressing ethical challenges. The New England journal of medicine, 378(11):981, 2018.

Laura L Colbran, Eric R Gamazon, Dan Zhou, Patrick Evans, Nancy J Cox, and John A Capra. Inferred divergent gene regulation in archaic hominins reveals potential phenotypic differences. Nature ecology & evolution, 3(11):1598–1606, 2019.

Ana Conesa, Pedro Madrigal, Sonia Tarazona, David Gomez-Cabrero, Alejandra Cervera, Andrew McPherson, Michał Wojciech Szcześniak, Daniel J Gaffney, Laura L Elo, Xuegong Zhang, et al. A survey of best practices for rna-seq data analysis. Genome biology, 17(1):13, 2016.

GTEx Consortium et al. The genotype-tissue expression (gtex) pilot analysis: multitissue gene regulation in humans. Science, 348(6235):648–660, 2015.

GTEx Consortium et al. Genetic effects on gene expression across human tissues. Nature, 550(7675): 204–213, 2017.

William Cookson, Liming Liang, Gonçalo Abecasis, Miriam Moffatt, and Mark Lathrop. Mapping complex disease traits with global gene expression. Nature Reviews Genetics, 10(3):184–194, 2009.

Valur Emilsson, Gudmar Thorleifsson, Bin Zhang, Amy S Leonardson, Florian Zink, Jun Zhu, Sonia Carlson, Agnar Helgason, G Bragi Walters, Steinunn Gunnarsdottir, et al. Genetics of gene expression and its effect on disease. Nature, 452(7186):423–428, 2008.

William E Evans and Mary V Relling. Moving towards individualized medicine with pharmacogenomics. Nature, 429(6990):464–468, 2004.

Eric R Gamazon, Heather E Wheeler, Kaanan P Shah, Sahar V Mozaffari, Keston Aquino-Michaels, Robert J Carroll, Anne E Eyler, Joshua C Denny, Dan L Nicolae, Nancy J Cox, et al. A gene-based association method for mapping traits using reference transcriptome data. Nature genetics, 47(9): 1091, 2015.

Eric R Gamazon, Ayellet V Segrè, Martijn van de Bunt, Xiaoquan Wen, Hualin S Xi, Farhad Hormozdiari, Halit Ongen, Anuar Konkashbaev, Eske M Derks, François Aguet, et al. Using an atlas of gene regulation across 44 human tissues to inform complex disease-and trait-associated variation. Nature genetics, 50(7):956–967, 2018.

Ian J. Goodfellow, Jean Pouget-Abadie, Mehdi Mirza, Bing Xu, David Warde-Farley, Sherjil Ozair, Aaron Courville, and Yoshua Bengio. Generative adversarial networks, 2014. Ahmed Hosny and Hugo JWL Aerts. Artificial intelligence for global health. Science, 366(6468): 955–956, 2019.

Haiyan Huang, Chun-Chi Liu, and Xianghong Jasmine Zhou. Bayesian approach to transforming public gene expression repositories into disease diagnosis databases. Proceedings of the National Academy of Sciences, 107(15):6823–6828, 2010.

Sergey Ioffe and Christian Szegedy. Batch normalization: Accelerating deep network training by reducing internal covariate shift. In Proceedings of the 32nd International Conference on International Conference on Machine Learning - Volume 37, ICML’15, page 448–456. JMLR.org, 2015.

Mary-Claire King and Allan C Wilson. Evolution at two levels in humans and chimpanzees. Science, 188(4184):107–116, 1975.

Roderick JA Little and Donald B Rubin. Statistical analysis with missing data, volume 793. John Wiley & Sons, 2019.

Leland McInnes, John Healy, Nathaniel Saul, and Lukas Grossberger. Umap: Uniform manifold approximation and projection. The Journal of Open Source Software, 3(29):861, 2018.

Tomas Mikolov, Ilya Sutskever, Kai Chen, Greg Corrado, and Jeffrey Dean. Distributed representations of words and phrases and their compositionality, 2013.

Cancer Genome Atlas Network et al. Comprehensive molecular portraits of human breast tumours. Nature, 490(7418):61, 2012.

Cancer Genome Atlas Research Network. Genomic and epigenomic landscapes of adult de novo acute myeloid leukemia. New England Journal of Medicine, 368(22):2059–2074, 2013.

Cancer Genome Atlas Research Network et al. Comprehensive molecular profiling of lung adenocarcinoma. Nature, 511(7511):543–550, 2014.

Marina Sirota, Joel T Dudley, Jeewon Kim, Annie P Chiang, Alex A Morgan, Alejandro Sweet-Cordero, Julien Sage, and Atul J Butte. Discovery and preclinical validation of drug indications using compendia of public gene expression data. Science translational medicine, 3(96):96ra77–96ra77, 2011.

Oliver Stegle, Leopold Parts, Matias Piipari, John Winn, and Richard Durbin. Using probabilistic estimation of expression residuals (peer) to obtain increased power and interpretability of gene expression analyses. Nature protocols, 7(3):500, 2012.

Daniel J Stekhoven and Peter Bühlmann. Missforest—non-parametric missing value imputation for mixed-type data. Bioinformatics, 28(1):112–118, 2012.

Xiaochen Sun, Santiago Vilar, and Nicholas P Tatonetti. High-throughput methods for combinatorial drug discovery. Science translational medicine, 5(205):205rv1–205rv1, 2013.

T. Tieleman and G. Hinton. Lecture 6.5—RmsProp: Divide the gradient by a running average of its recent magnitude. COURSERA: Neural Networks for Machine Learning, 2012.

Olga Troyanskaya, Michael Cantor, Gavin Sherlock, Pat Brown, Trevor Hastie, Robert Tibshirani, David Botstein, and Russ B Altman. Missing value estimation methods for dna microarrays. Bioinformatics, 17(6):520–525, 2001.

Michael E Wall, Andreas Rechtsteiner, and Luis M Rocha. Singular value decomposition and principal component analysis. In A practical approach to microarray data analysis, pages 91–109. Springer, 2003.

John N Weinstein, Eric A Collisson, Gordon B Mills, Kenna R Mills Shaw, Brad A Ozenberger, Kyle Ellrott, Ilya Shmulevich, Chris Sander, Joshua M Stuart, Cancer Genome Atlas Research Network, et al. The cancer genome atlas pan-cancer analysis project. Nature genetics, 45(10):1113, 2013a.

John N Weinstein, Eric A Collisson, Gordon B Mills, Kenna R Mills Shaw, Brad A Ozenberger, Kyle Ellrott, Ilya Shmulevich, Chris Sander, Joshua M Stuart, Cancer Genome Atlas Research Network, et al. The cancer genome atlas pan-cancer analysis project. Nature genetics, 45(10):1113, 2013b.

Genevieve L Wojcik, Mariaelisa Graff, Katherine K Nishimura, Ran Tao, Jeffrey Haessler, Christopher R Gignoux, Heather M Highland, Yesha M Patel, Elena P Sorokin, Christy L Avery, et al. Genetic analyses of diverse populations improves discovery for complex traits. Nature, 570(7762): 514–518, 2019.

Jinsung Yoon, James Jordon, and Mihaela Van Der Schaar. Gain: Missing data imputation using generative adversarial nets. arXivpreprint arXiv:1806.02920, 2018.

Bin Zhang and Steve Horvath. A general framework for weighted gene co-expression network analysis. Statistical applications in genetics and molecular biology, 4(1), 2005.

Øystein Sørensen, Kristoffer Herland Hellton, Arnoldo Frigessi, and Magne Thoresen. Covariate selection in high-dimensional generalized linear models with measurement error. Journal of Computational and Graphical Statistics, 27(4):739–749, 2018. doi: 10.1080/10618600.2018.1425626. URL https://doi.org/10.1080/10618600.2018.1425626.

